# Deconvoluting Virome-Wide Antiviral Antibody Profiling Data

**DOI:** 10.1101/333625

**Authors:** Daniel R. Monaco, Sanjay V. Kottapalli, Tiezheng Yuan, Florian P. Breitwieser, Danielle E. Anderson, Limin Wijaya, Kevin Tan, Wan Ni Chia, Kai Kammers, Mario Caturegli, Kathleen Waugh, Marian Rewers, Lin-Fa Wang, H. Benjamin Larman

## Abstract

The ability to comprehensively characterize exposures and immune responses to viral infections will be critical to better understanding human health and disease. We previously described the VirScan system, a phage-display based technology for profiling antibody binding to a comprehensive library of peptides designed to represent the human virome. The previous VirScan analytical approach did not fully account for disproportionate representation of viruses in the library or for antibody cross-reactivity among sequences shared by related viruses. Here we present the ‘AntiViral Antibody Response Deconvolution Algorithm’ (‘AVARDA’), a multi-module software package for analyzing VirScan datasets. AVARDA provides a probabilistic assessment of infection at species-level resolution by considering alignment of all library peptides to each other and to all human viruses. We employed AVARDA to analyze VirScan data from a cohort of encephalitis patients with either known viral infections or undiagnosed etiologies. By comparing acute and convalescent sera, AVARDA successfully confirmed or detected antibody responses to human herpesviruses 1, 3, 4, 5, and 6, thereby improving the rate of diagnosing viral encephalitis in this cohort by 62.5%. We further assessed AVARDA’s utility in the setting of an epidemiological study, demonstrating its ability to determine infections acquired in a child followed prospectively from infancy. We consider ways in which AVARDA’s conceptual framework may be further developed in the future and describe how its analyses may be extended beyond investigations of viral infection. AVARDA, in combination with VirScan and other pan-pathogen serological techniques, is likely to find broad utility in the epidemiology and diagnosis of infectious diseases.

## Introduction

Comprehensive profiling of virome-wide antibody binding specificities has broad utility for epidemiological investigations, surveillance of emerging viruses, and the unbiased diagnosis of infections^1,2,3,4^. Phage ImmunoPrecipitation Sequencing (“PhIP-seq”)^5^ with a peptide library spanning the human virome (“VirScan”)^6^ provides a platform for unbiased, high-throughput, low cost analysis of anti-viral antibodies. While other multiplexed serological techniques exist^7^, each is limited in its representation of viral antigens^8^, the size and quality of the epitopes presented^9^, the per-sample assay cost and/or sample throughput. VirScan provides favorable performance characteristics, but until now interpretation of assay results has been limited by a rudimentary analytical framework.

Our previously published analytical approach suffers from three critical limitations. First, the number of non-overlapping, virus-associated antibody specificities (a measure of response “breadth”) conveys important biological information and informs the confidence of a predicted exposure. Non-overlapping specificities were defined using a suboptimal heuristic that typically underestimated response breadth. Second, a significantly enriched peptide was considered only in the context of the specific virus it was designed to represent. This ignored sequence homology between related viruses, and any potential for antibody cross-reactivity. Further, the VirScan library was designed to cover single representative proteins from UniProt clusters of 90% identity. Relying solely upon the intended viral representations of enriched peptides to diagnose infections will therefore result in both false negative results (“missing” peptides from the true infective organism) and false positive results (peptide enrichments due to cross-reactive antibodies). Third, we previously relied on each virus’s proteome “size” to establish virus-specific thresholds for seropositivity. This approach ignored the proportional representation of each virus within the enriched set of peptides and the corresponding virus-specific representation in the library. Additionally, using a threshold for seropositivity is far less informative than a probabilistic assessment of a viral infection.

Here we introduce the <u>A</u>nti-<u>V</u>iral <u>A</u>ntibody <u>R</u>esponse <u>D</u>econvolution <u>A</u>lgorithm (‘AVARDA’), an analytical framework for improved analysis of massively multiplexed anti-viral antibody binding data. AVARDA integrates three key modules to generate a conservative and probabilistic assessment of current and historical viral exposures. The first module employs a network graph based on peptide-peptide relationships to define the minimum number of independent antibody specificities (i.e. response breadth) required to completely explain the set of observed enriched peptides. The second module uses sequence alignment to define each peptide’s relationship to a comprehensive database of all human viral genomes, which have been translated in all six reading frames. This permits a conservative consideration of *all* viral infections that could have *potentially* elicited the observed antibodies specificities, and conversely, which *sets* of peptide enrichments could *potentially* be associated with each virus. The third module iteratively assigns each peptide to its most likely associated viral infection(s), according to a null model that considers the representation of each virus in the VirScan library. Our approach permits peptides to be associated with multiple distinct infections, provided there is sufficient evidence for each viral infection on its own (i.e. in the absence of the shared peptides). AVARDA also indicates uncertainty in peptide-virus assignments when there is insufficient evidence to discriminate between infections by related viral candidates, but when there is enough evidence to conclude that an infection has indeed occurred. Linking these modules, the final output of AVARDA provides multiple hypothesis adjusted p-values for exposure to each virus, along with the associated breadths of the antibody responses and the relationships between the enriched peptides.

The utility of antibody-based diagnostics has been constrained by an inability to robustly discriminate among infections by closely related viral pathogens^10^. An important example is the challenge of distinguishing between flavivirus infections, notably in the setting of the recent Zika virus outbreak^11^. Unbiased analysis of anti-viral antibodies using a robust algorithm that comprehensively considers potential cross-reactivity will dramatically improve the clinical utility of antibody analyses. Single time-point testing of IgG reactivity is difficult to interpret in the context of an ongoing or recent infection, as these antibodies may be due to long lived plasma cells induced by a historic infection. Quantifying an individual’s *change* in IgG specificities over time, however, may provide clearer evidence of an evolving immune response. Similarly, comparing IgG specificities across distinct physiological compartments may provide evidence of an organ-specific infection (e.g., blood versus cerebrospinal fluid in the setting of encephalitis). Such pairwise VirScan comparisons would be particularly problematic for our previous analytical approach, because (i) a low number (but high proportion) of changing peptides may be observed during response to a single infection, and (ii) ongoing antibody responses may be more associated with broadly cross-reactive antibodies^12^. We have therefore developed the AVARDA algorithm using pairwise VirScan data generated longitudinally from patients with viral encephalitis. The predictions of the algorithm were found to agree with clinical nucleic acid or serology testing in most cases. In addition, utilizing VirScan/AVARDA analysis in conjunction with traditional nucleic acid testing resulted in a 62.5% increase in the diagnosis of viral encephalitis. Finally, we demonstrate the utility of AVARDA for epidemiological investigations by determining childhood infections acquired in the setting of a longitudinal cohort study.

## Results

### Differential quantification of antibody specificities

We performed VirScan analysis on paired sera obtained from an encephalic cohort, collected upon initial hospital admission and fourteen days post-admission. To quantify changes in their anti-viral antibody repertoire, we adapted our recently reported z-score based algorithm to the pairwise differential quantification of peptide enrichments^13^. Changes in antibody binding over time were considered significant when the initial versus day 14 pairwise z-scores were greater than 10. **Fig 1A** shows the results of one patient’s (Pt1) pairwise VirScan analysis. All pairwise enriched peptides and the viral species they were intended to represent are provided in Supplemental Table 1. Nucleic acid testing had previously diagnosed a herpes simplex virus 1 (HSV1) infection in this individual, consistent with the predominance of enriched peptides designed to represent alphaherpesvirus-associated proteins. Notably, antibodies from this patient with HSV1 encephalitis enriched 22 peptides designed to represent HSV2, highlighting the potential cross-reactivity of anti-viral antibodies.

**Table 1.**
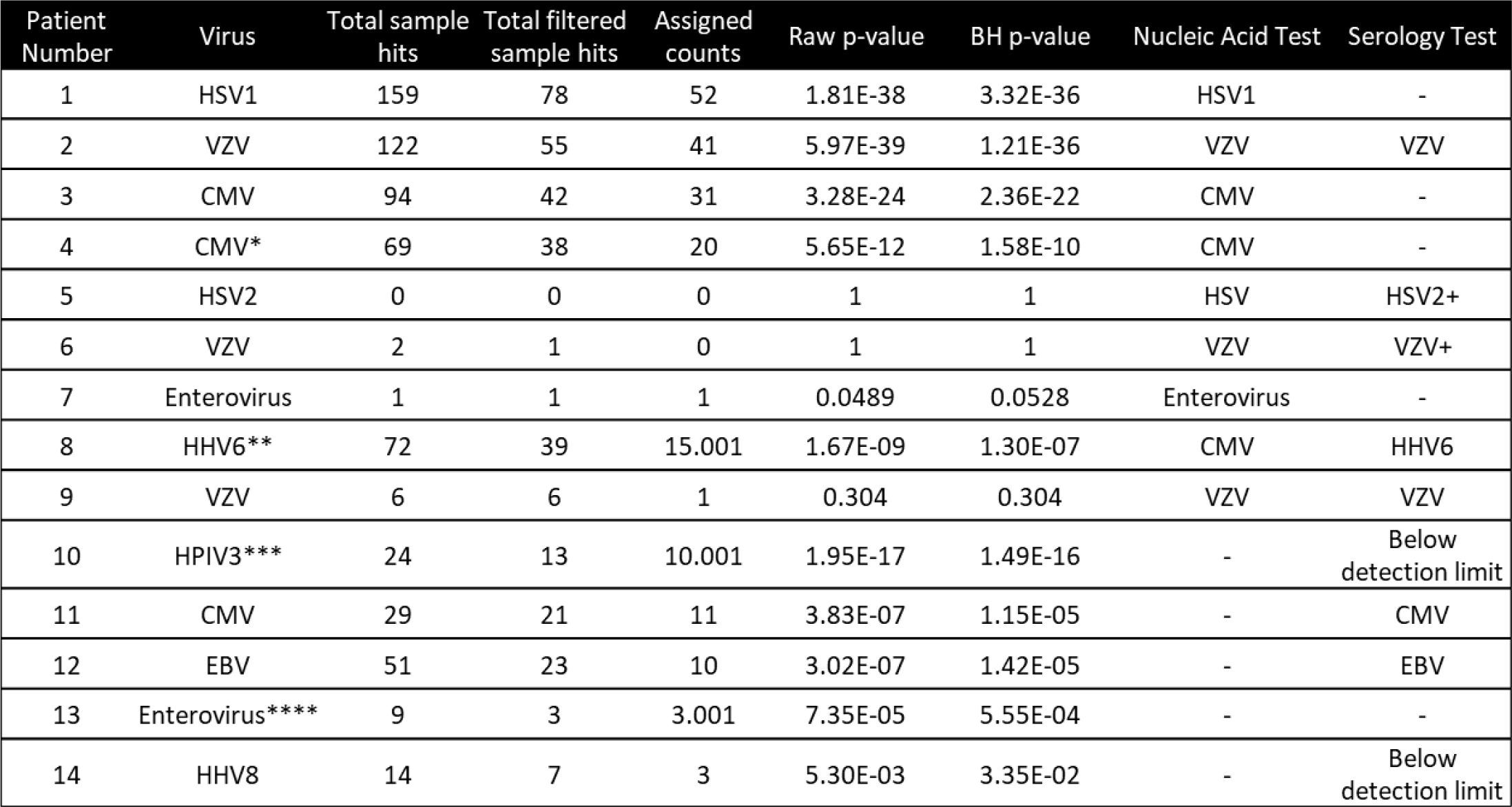
Summary of AVARDA results from the encephalitis cohort. The ‘virus’ column indicates the identified viral infection via AVARDA or nucleic acid test. The ‘total sample hits’ and ‘total filtered sample hits’ refer to the number of total pairwise enriched peptides and the number of enriched peptides after independence filtering by Module 1. The ‘assigned counts’ indicates the number of enriched peptides associated with the significant infection after AVARDA analyses. The ‘Raw p-value’ and the ‘BH p-value’ columns represent original and multiple hypothesis test corrected AVARDA significance values respectively. Nucleic acid or serologic test results are indicted in the last two columns. ‘–’ indicates that a specific test was not conducted. * AVARDA also identified an EBV infection with a pBH = .046. **HHV6A and B infections were both indistinguishably identified but reported as HHV6. ***HPIV shared an indistinguishability tag with bovine respirovirus. **** Pt13’s enterovirus infection shared indistinguishability tags with enteroviruses A – C, H and Rhinovirus A – C. + indicates serology testing that did detect virus specific IgG but it was unchanged between timepoints.

**Figure 1.**
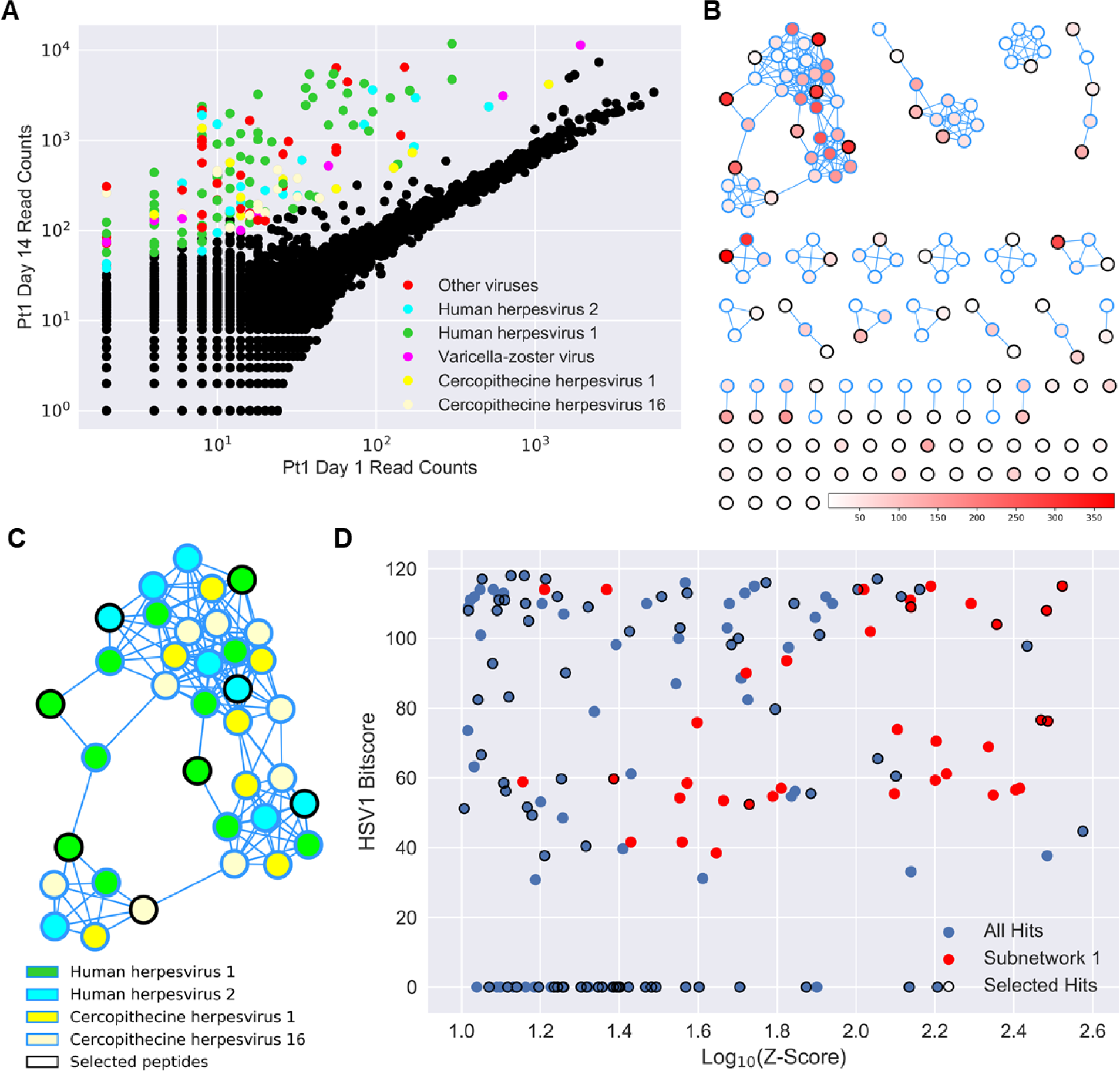
Cross reactivity of an antibody response. **(A)** VirScan normalized read counts plotted for Pt1 at day 14 versus day 1 (hospital admission). Peptides of pairwise z-score > 10 are colored according to the virus they were designed to represent. Network graph representation of peptide overlaps within the enriched set of peptides shown in A. Nodes represent peptides and edges are drawn between peptides that share at least one 7-amino acid stretch of identity. Peptides are colored according to z-score and outlined in black if selected by the independence filter. **(C)** Peptides within the largest connected subgraph are colored according to the virus they were designed to represent. **(D)** BLAST score of each peptide from A aligned against HSV1, plotted against the log10 of their pairwise z-scores. Peptides are outlined in black if they were selected by the independence filter.

### Module 1: Defining the maximal set of nonoverlapping antibody specificities

The VirScan library was designed to represent the proteins of the human virome as 56 amino acid peptide tiles, with adjacent tiles overlapped by 28 amino acids. Due to these designed overlaps, as well as the presence of homologous peptides from related viruses, a wide range of epitope redundancy exists within the library. As a simple preliminary model, we assumed that two peptides sharing an exact sequence at least seven amino acids long could potentially be recognized by the same monoclonal antibody. Conversely, if two peptides were enriched and did not share a seven amino acid sequence, we assumed that their binding was due to at least two unique antibody clones. We therefore constructed a database of all seven amino acid overlaps among all peptides in the VirScan library. For any set of enriched peptides, this database can be used to construct a network graph, such that peptides are represented by nodes and overlapping sequences are represented by edges connecting the nodes. **Fig 1B** illustrates the network of Pt1’s pairwise-enriched peptides.

Given a network of enriched peptides, we devised an iterative method to remove peptides until a maximal subset of unconnected peptides remains. This number of remaining peptides represents the minimum number of non-redundant antibody specificities required to produce the network, and can therefore be considered a conservative estimate of the antibody response breadth. In order to iteratively disconnect the graph, we remove the most connected peptide(s) one at a time. Ties are broken by favoring retention of more strongly enriched peptides, based on the assumption that these peptides are more likely to harbor cognate, versus cross reactive epitopes. This process is further described in our methods. The final set of non-overlapping (“independent”) peptides selected from Pt1’s network using this approach are denoted with solid outlines in **Fig 1B**. We also examined Pt1’s largest connected subgraph and the viruses which these peptides were designed to represent (**Fig 1C**).

To assess the potential for unintended selection bias in our graph disconnection approach (such as inadvertantly favoring lower alignment scores or weaker enrichments), we performed two Mann-Whitney U tests to detect significant differences in z-scores or HSV1 alignment scores between the set of selected non-overlapping peptides versus the set of peptides discarded during network disconnection. HSV1 alignment scores are plotted versus log10 z-scores in **Fig 1D**. Enriched peptides with zero connections were not considered in this analysis as they are unaffected by our iterative method. These tests did not reach significance (two-tailed p-values of 0.19 and 0.46, respectively), suggesting that this method of non-overlapping peptide selection does not significantly bias the z-scores or alignment scores of the selected peptides. From Pt1’s 159 pairwise-enriched peptides, 78 non-overlapping peptides were selected in this way for further analysis.

### Module 2: Enumerating all possible peptide-virus associations

In our previous study, each VirScan peptide had been defined simply by the specific viral protein it was intended to represent. However, in order to exhaustively consider sequence similarity and thus potential for antibody cross reactivity, we have constructed a comprehensive genomic database of all known human viral species. To exclude any potential open reading frame (ORF) annotation bias, all VirScan peptide sequences were aligned via tblastn against a protein sequence database formed by translating these viral genomes in all 6 reading frames^14^. The relationship of all VirScan peptides to all human viruses can thus be represented as a matrix of alignment bit scores. **Fig 2A** contrasts the design of Pt1’s non-overlapping enriched peptides, versus their alignment in this matrix. The numbers of these peptides which aligned to only one single virus are also reported. Note that consideration of peptide-virus alignments enables the possibility to detect infections involving viruses not explicitly represented in the VirScan library design (e.g., Macacine alphaherpesvirus 1). As the reference database is updated, it is thus possible that new infections may be identified by re-analyzing existing data. The matrix of alignment bitscores associated with Pt1’s enriched, non-overlapping peptides is provided as a clustered heatmap in **Fig 2B**.

**Figure 2.**
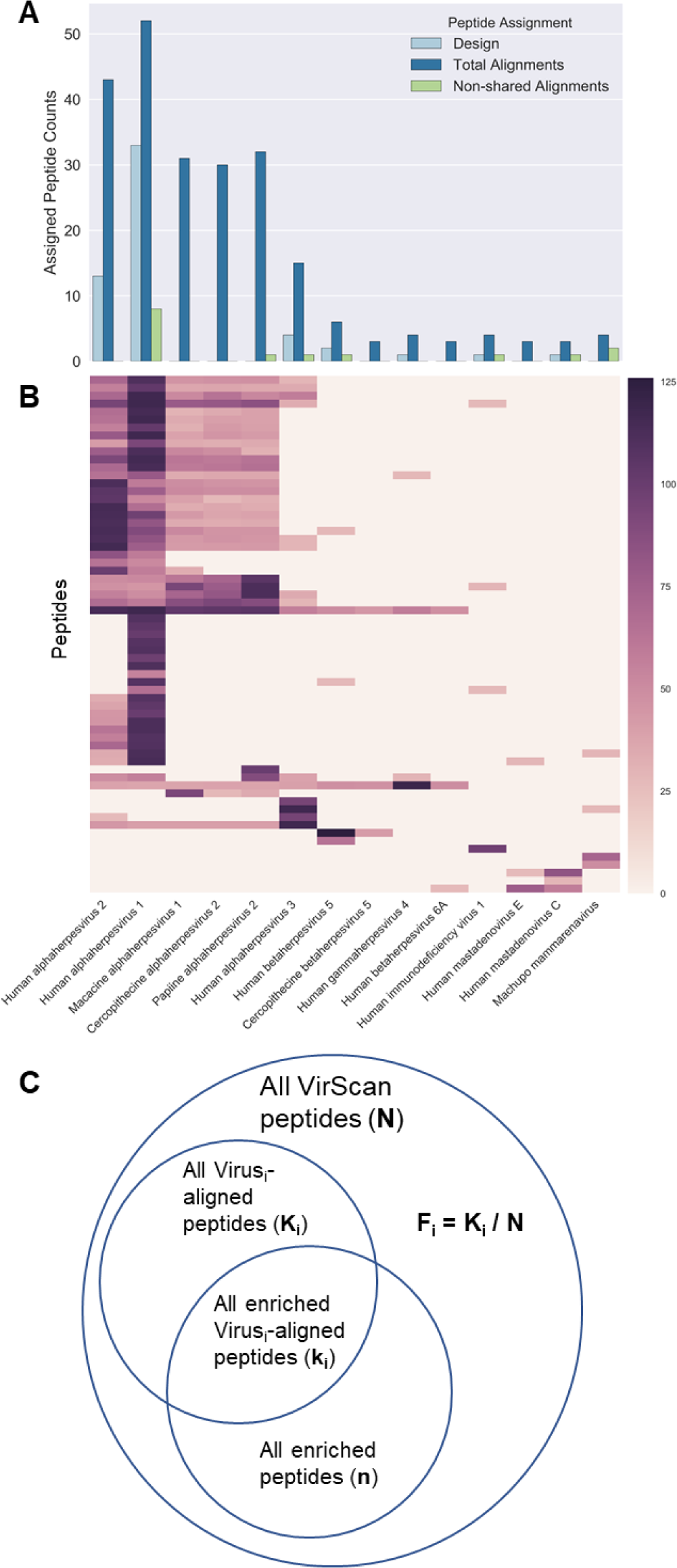
Associating peptides and viruses using alignment. **(A)** Number of Pt1’s enriched peptides associated with the most significant viruses. Numbers of enriched peptides designed to represent each virus are shown in light blue. Total numbers of enriched peptide alignments to each virus are shown in dark blue. The peptide alignments not shared with any other viruses are shown in green. **(B)** Clustered heatmap of the enriched peptide alignments, colored by bitscore of blast alignment. **(C)** Diagram depicting the calculation of the binomial null probability for one virus, given a set of enriched peptides.

**Fig 2B** illustrates that most of Pt1’s enriched, non-overlapping peptides align with multiple viruses. Also apparent is that most of these alignments are shared among the human alphaherpesviruses. As expected, given the diagnosed HSV1 infection, however, almost every single peptide that aligns with an alphaherpesvirus, aligns with HSV1. Using a binomial test, we then calculated the probability that the fraction of HSV1-aligning selected peptides among all selected peptides would occur under a null model defined by the fractional representation of HSV1-aligning peptides in the entire VirScan library (**Fig 2C**). In this way, Module 2 is used to rank order all potential infections by significance in preparation for the next stage of analysis.

### Module 3: Conservatively assessing potential infections

Implementing Module 2 results in increased sensitivity and decreased specificity for detecting infections, by associating all independently enriched peptides with all possible viral infections. The goal of Module 3 is to weigh the evidence for each infection on its own, given the relationship of all its enriched peptides to all other viruses. Initially, the most significant virus (‘virusi’), as determined by Module 2, is iteratively compared with viruses that also share peptide alignments (‘virusj’). The numbers of shared and non-shared peptides are separately evaluated for significance, using binomial testing as described in the Methods.

Three possible outcomes for each virusi-virusj comparison are illustrated in **Fig 3A**: (i) The number of non-shared virusi-associated peptides is significant, while the number of non-shared virusj-associated peptides is not significant, suggesting that enrichment of the shared peptides can likely be attributed to infection *only* with virusi. In this case, peptides aligning to both viruses are no longer associated with virusj. (ii) The number of non-shared peptides are significant for both viruses, suggesting that the uniquely aligning and the shared enrichments can be attributed to infections by *both* virusi and virusj. In this case, the shared peptides remain associated with both viruses. (iii) The number of shared peptides is significant, but the number of non-shared peptides associated with either virus alone is not significant. In this scenario we assume the enrichments are most likely due to a true infection, but that there is insufficient evidence to distinguish whether an infection originated from *either* or *both* viruses. In this case the shared peptide alignments remain shared, but both viruses are assigned a tag to indicate that their infections could not be distinguished. After each iteration of this probabilistic assessment of shared peptide-virus associations, enriched peptides associated exclusively with virusi are removed from consideration. An updated p-value is then calculated for each remaining virus, and the process is iterated until all virus pairs with shared peptide alignments have been analyzed. Upon completion, Module 3 will have therefore conservatively removed false positive viral associations, while having retained each peptide’s association with the most likely causative infection(s), including indistinguishable infections.

**Figure 3.**
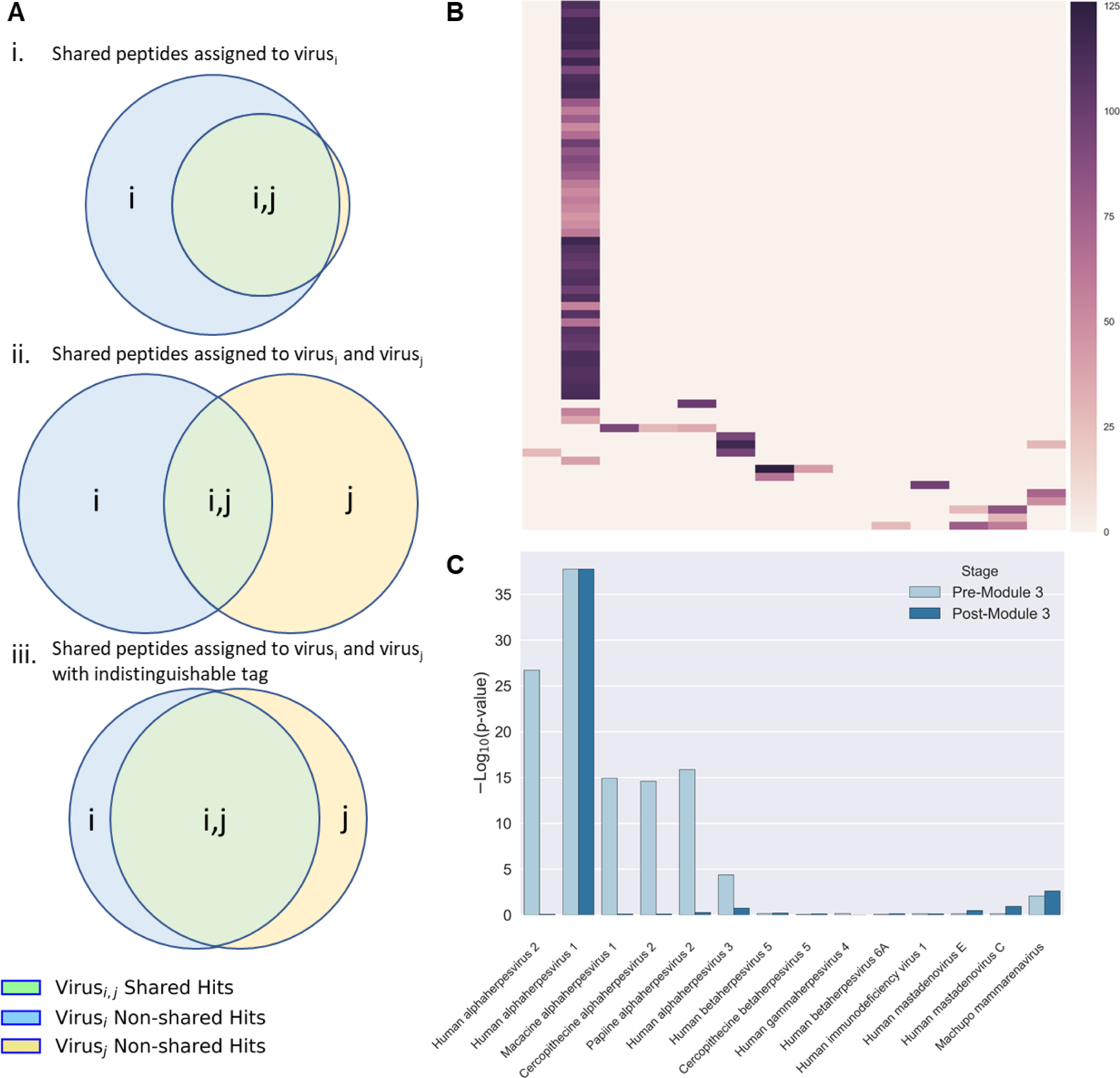
Treatment of enriched peptides with alignments to multiple viruses **(A)** Schematic representing three modes of shared peptide alignments between two viruses: (i) single infection by virusi can account for all alignments to virusj, (ii) infections caused by both virusi and virusj, and (iii) an infection by virusi and/or virusj which cannot be distinguished. **(B)** The peptide alignment heatmap of Figure 2B after removal of alignments determined to be cross-reactive. **(C)** Bar graph of-log10(p-values) for each virus before and after consideration of cross-reactivity using Module 3.

**Fig 3B** shows the alignment matrix of **Fig 2B** updated after Module 3’s operations upon the shared peptide alignments. The final binomial test-generated p-value for each virus is then adjusted for multiple hypothesis testing using the Benjamini-Hochberg (“BH”) procedure^15^. In this way, Pt1’s data yielded a single significant infection, HSV1, with a significance of pBH = 3×10^−36^. A direct comparison of the pre- and post-Module 3 p-values is provided in **Fig 3C**. Module 3 generates several intermediate files containing useful information for understanding how each peptide was evaluated during the course of its analysis.

### AVARDA for unbiased diagnosis of viral encephalitis

The AVARDA framework used to analyze Pt1’s longitudinal data was applied to an additional set of 37 encephalitic patients with potential viral etiologies. Of these, eight had received diagnoses of viral infection based on clinical nucleic acid testing; the remainder were undiagnosed and treated as potential viral encephalitis. Summarized findings are provided in Table 1. Using AVARDA with thresholds requiring at least three independent enriched peptides and a pBH < 0.05, we identified statistically significant extant viral infections in nine of these patients. The pairwise VirScan data sets are provided as **Fig 4**. A similar AVARDA analysis of 38 encephalitic patients not suspected of having viral encephalitis resulted in only a single significant infection (Pt14). Interestingly, this infection barely exceeded threshold with three aligning independent peptides and a relatively weak pBH = .03. Four of the nine cases with diagnoses confirmed by nucleic acid testing (Pt1, Pt2, Pt3, and Pt4) corresponded precisely with the infections indicated by AVARDA. Results for Pt8 were discordant, however, with a PCR based diagnosis of cytomegalovirus (CMV), versus AVARDA’s determination of roseolavirus infection (X independent HHV6 peptide enrichments, pBH = 1.3×10^−7^). Closer investigation of Pt8’s clinical history revealed previous Ganciclovir treatment for CMV infection, suggesting a long term prior infection. Consistent with this, AVARDA detected significant CMV antibodies at both day 1 and 14, but their reactivity was differentially unchanged over time. Clinical antibody testing agreed with the results of AVARDA, measuring increased HHV6-specific IgG in the convalescent serum. The remaining four patients were diagnosed with HSV (Pt5), enterovirus (Pt7), and varicella-zoster virus (VZV, Pt6 and Pt9, **Fig 4E** and **Fig S1**, respectively), but none had significant pairwise changes detected by AVARDA over the two week period (**Fig 4**).

**Figure 4.**
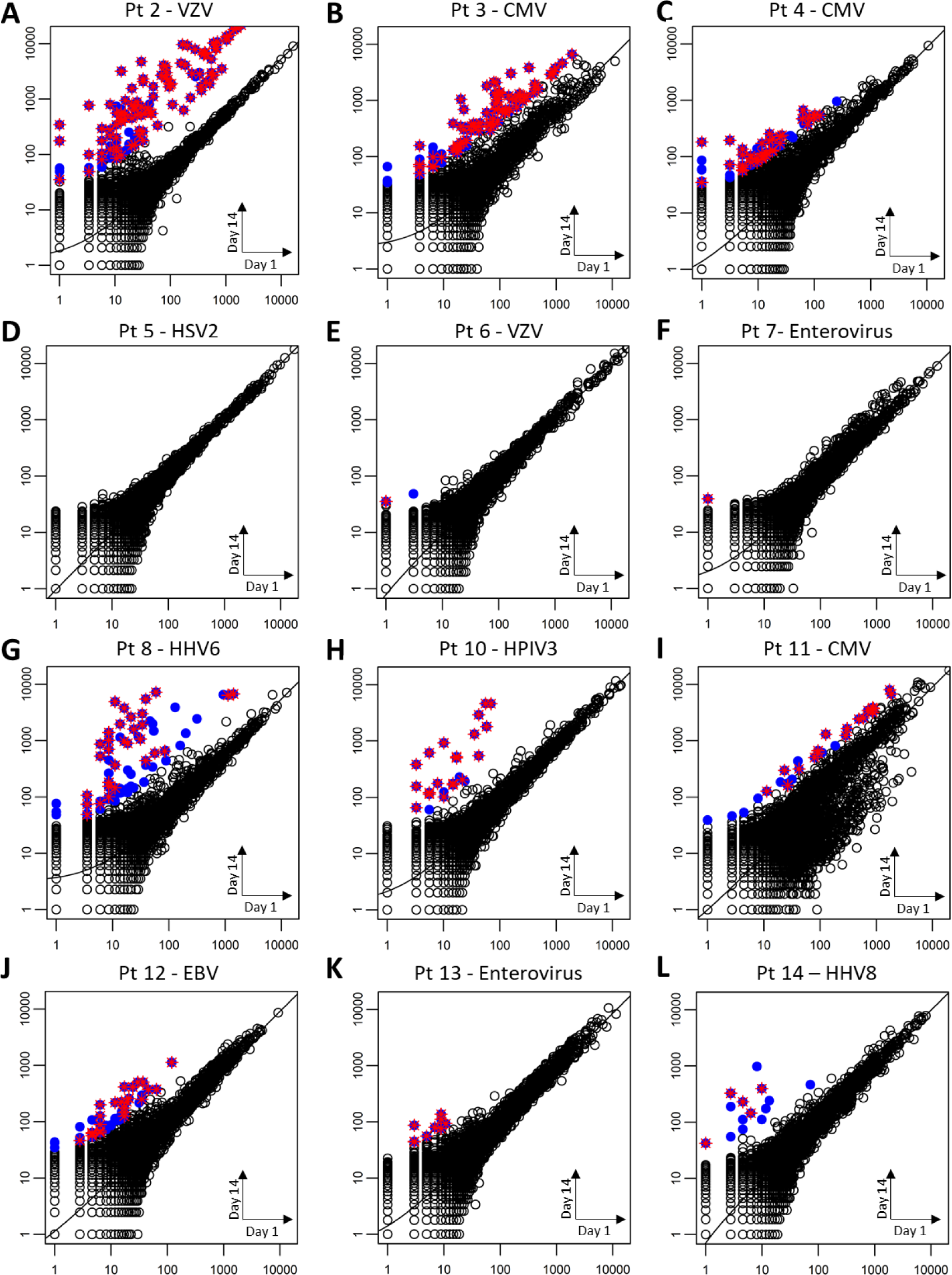
AVARDA analysis of pairwise enrichment scores from twelve patients with encephalitis. Scatterplots **A-L** correspond to patients 2-13 in Table 1. Each plot indicates the VirScan normalized read counts from the day 1 dataset (x- axis) plotted against the day 14 dataset (y-axis). Robust linear regression lines are drawn for each scatterplot. Points in blue represent peptides enriched with pairwise z-scores >10. Enriched peptides associated with a significant infection determined by AVARDA are additionally marked with a red cross. Pt9’s scatterplot is provided in Fig S1.

Single timepoint AVARDA analyses and clinical antibody testing both detected high but unchanging virus specific IgG levels in Pt5 and Pt6. This suggests that either (i) these were chronic, pre-existing infections potentially unrelated to their encephalitis, that (ii) these patients’ initial sera were collected too late in the course of their disease to see differential changes, or that (iii) VirScan did not detect the changing antibody specificities/titers. Clinical antibody testing detected increasing VZV-specific IgG levels in Pt9, which was not detected by AVARDA. Further investigation of Pt9’s VirScan results, however, revealed several VZV aligning peptides that appeared to be differentially enriched, but below our threshold, indicating that our default threshold may be too stringent in some cases (**Fig S1**). In 4 individuals with undiagnosed encephalitis, AVARDA identified significant infections with HPIV3 (Pt10), CMV (Pt11), Epstein-Barr Virus (EBV, Pt12) and enterovirus (Pt13). Confirmatory clinical serology detected increasing viral IgG for two of these cases (CMV for Pt11 and EBV for Pt12). We could not confirm Pt10’s AVARDA findings, however, potentially due to limited availability of serum, which was diluted prior to testing. Additionally, we were unable to conduct clinical serology tests for enterovirus IgG (for Pt7 and Pt13), due to insufficient serum volume.

### Use of AVARDA for longitudinal studies

The pathophysiological sequellae of infectious diseases are believed to contribute to many human illnesses, including autoimmunity (e.g., type 1 diabetes)^16^, neurodegenerative disease (e.g., Alzheimer’s disease)^17^, cancer (e.g., cervical cancer)^18^ and allergic disease (e.g., asthma)^19^. Longitudinal cohort studies provide important epidemiological approaches for causally linking environmental exposures, such as viral infection, with (pre-)clinical outcomes. To assess the utility of VirScan/AVARDA in this setting, we profiled serum samples collected longitudinally every two to six months at-risk individuals, prior to the onset of type 1 diabetes. **Table 2** summarizes the results of AVARDA’s analyses for one study participant across 14 available time points from 9 months to 5 years of age. AVARDA detected 20 infections (including responses to immunizations) that met our significance criteria over this time course (**Fig 5**). These included adenovirus, enterovirus, parechovirus, rhinovirus, influenza, norovirus, parainfluenza virus, metapneumovirus, respiratory syncytial virus, coronavirus, and astrovirus infections. Antibodies directed against measles and rubella, but not mumps, were detected in the first interval after the MMR vaccine was administered at 1.05 years of age. **Fig 5** also highlights the longitudinal behavior of 23 peptides from parechovirus, which were found to be differentially enriched by comparison of sera obtained at 1.67 versus 1.26 years of age. This type of peptide-level analysis can be used to characterize the epitope specific temporal antibody responses, a subset of which may persist due to the formation of long lived plasma cells.

**Table 2.** AVARDA results from one individual’s longitudinally collected sera. The ‘virus’ column indicates the viral infection identified by AVARDA. The ‘Age Interval(s) of Infection’ column refers to the patient age interval in which a given infection was identified; if the same infection was identified at two consecutive time intervals it was considered the same infection and the intervals of infection were merged. The ‘total sample hits’ and ‘total filtered sample hits’ columns provide the number of total pairwise enriched peptides and the number of enriched peptides after implementation of Module The ‘assigned counts’ indicates the number of enriched peptides associated with the significant infection after AVARDA analyses. The ‘Raw p-value’ and the ‘BH p-value’ columns represent original and multiple hypothesis test corrected AVARDA significance values respectively.

| Virus | Age Interval(s) of Infection (years) | Total sample hits | Total filtered sample hits | Assigned counts | Raw p-value | BH p-value |
| --- | --- | --- | --- | --- | --- | --- |
| Mastadenovirus | 0.76 to 1.67 | 323 | 72 | 33 | 1.18E-24 | 1.40E-22 |
|  | 3.29 to 3.58 | 90 | 48 | 10 | 7.24E-05 | 6.77E-03 |
| Measles | 0.76 to 1.26 | 323 | 72 | 5.002 | 1.71E-04 | 2.27E-03 |
| Rubella | 0.76 to 1.26 | 323 | 72 | 3 | 2.45E-03 | 2.08E-02 |
| Enterovirus | 0.76 to 1.26 | 323 | 72 | 19.001 | 1.75E-14 | 1.04E-12 |
|  | 1.67 to 2.15 | 73 | 11 | 5.001 | 2.52E-06 | 5.65E-05 |
|  | 3.83 to 4.43 | 136 | 16 | 10.001 | 1.38E-10 | 1.28E-09 |
|  | 4.73 to 5 | 80 | 41 | 8.001 | 1.51E-04 | 7.20E-03 |
| Parenchovirus | 1.26 to 1.67 | 77 | 32 | 7 | 2.06E-07 | 2.39E-05 |
| Influenza C | 1.67 to 1.9 | 73 | 11 | 4 | 3.95E-05 | 3.04E-04 |
| Norovirus | 2.15 to 2.46 | 40 | 23 | 5.001 | 6.04E-05 | 8.69E-03 |
| RSV | 2.46 to 2.97 | 104 | 28 | 5.001 | 4.97E-05 | 1.01E-02 |
| Metapneumovirus | 3.29 to 3.58 | 90 | 48 | 4 | 7.07E-05 | 6.67E-03 |
|  | 4.73 to 5 | 80 | 41 | 8 | 1.01E-10 | 1.20E-08 |
| Astrovirus | 4.43 to 5 | 66 | 19 | 3.002 | 2.20E-04 | 4.34E-03 |

**Figure 5.**
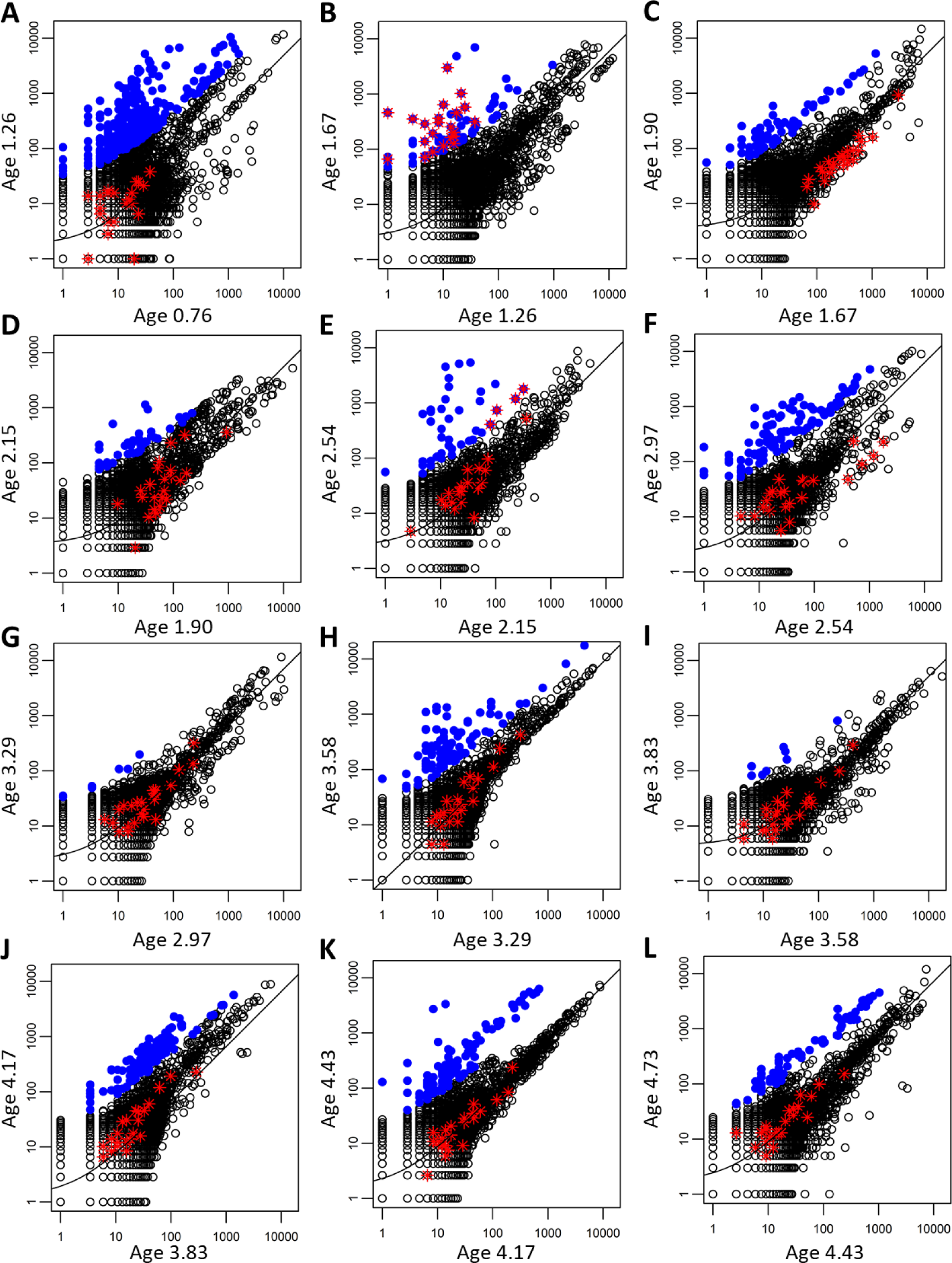
AVARDA analysis of pairwise enrichment scores from one patient with sera collected longitudinally. Scatterplots **A-L** corresponds to consecutive pairwise time intervals from 9 months to 5 years of age. Significant infections identified by AVARDA are detailed in Table 2. Robust linear regression lines are drawn for each scatterplot. Points in blue represent peptides enriched with pairwise z-scores >10. Enriched peptides associated with a parechovirus infection determined by AVARDA in **B** are additionally marked with a red cross in all plots.

Sensitivity for detecting infections may be diminished by infrequent sampling, due (i) to the transient nature of some humoral responses, (ii) to increased level inter-relationships among the differentially enriched set of peptides or (iii) to ‘lumping’ of sequential infections by related viruses. We therefore assessed the impact of sampling frequency on infection detection sensitivity. By selectively performing pairwise analyses across different sampling intervals, we observed the number of detected infections to decay with decreased sampling frequency (**Fig S2**). The design of longitudinal studies that include comprehensive antibody analyses should take these findings into consideration.

## Discussion

The diagnostic and epidemiologic utility of antibody analyses has been limited in large part due to the difficulty in accounting for antibody cross reactivities among closely related organisms^10^. The development of an analytical framework that accounts for potential cross reactivity in pan-viral serological data, and which provides probabilistic assessments of infections, is therefore an important advance. Our current study has demonstrated that AVARDA, when applied to VirScan analysis of longitudinally collected serum samples, can reliably identify evolving IgG responses against specific viruses.

We have examined the performance of AVARDA in diagnosing viral encephalitis, a setting in which it is often challenging, yet critical, to distinguish infectious from autoimmune etiologies^20,21^. As such, pan-viral IgG analysis may provide information complementary to nucleic acid testing, thus improving diagnostic sensitivity for low abundance or unexpected viruses, and for distinguishing active from latent infections. In this study, combining VirScan/AVARDA analysis with traditional diagnostic methods increased the rate of diagnosis by 62.5%. Pan-viral IgM analyses, while not explored here, may also be informative in differentiating new from recurrent exposures. Despite expanding the utility of viral antibody testing, VirScan/AVARDA is still limited in that it requires the development of an adaptive immune response, which takes valuable time and may not occur in patients with suppressed immune systems.

We propose four general areas in which future iterations of AVARDA could more effectively incorporate VirScan information. First, peptides enriched above a threshold are considered binary hits after Module 1, since their degrees of enrichment are not considered beyond the network graph filtering stage. If the contribution of each peptide to the final results were to be somehow weighted according to the magnitude of its enrichment, the confidence of the associated infection(s) could potentially be calculated more accurately. Second, the strengths of peptide-virus alignments, quantified as bit-scores, are similarly not considered by AVARDA. Incorporating these quantitative measures might also improve the final results. Third, enriched sets of peptides sharing a seven-amino acid sequence are currently treated as a single antibody specificity. However, the confidence associated with the enrichment of overlapping peptides is in general higher than for peptides enriched in isolation. If Module 1 attributed greater confidence to these peptide clusters, the sensitivity of AVARDA may be increased, particularly for infections eliciting more restricted responses. Additionally, AVARDA may benefit from a more nuanced consideration of overlapping peptides, versus the current simple requirement for a 7-aa perfect match overlap. Fourth, it may be possible to develop a statistical test that can distinguish among infections which are currently found to be indistinguishable by AVARDA. One can envision pairwise comparisons of homologous peptides that might discern the more likely of two highly related species (or even strains), when there is a consistent preference for antibody recognition of one organism’s peptides over those of another. If such an approach were successful, it would enable expanding the taxonomic reference database beyond the virus species level, potentially enhancing the specificity of AVARDA. Even at the species level, AVARDA frequently cannot distinguish highly interrelated viruses, particularly enteroviruses and adenoviruses in our experience. We expect that these, and other improvements not contemplated here, will markedly improve the performance of future versions of AVARDA.

We investigated the potential loss of sensitivity due to network graph filtering in Module 1, by executing AVARDA without invoking this step. Pt17, previously with no AVARDA-detected infections, exhibited nine increasing rubella virus peptide enrichments and six increasing herpesvirus 8 peptide enrichments (**Fig S3**). Clinical serological analysis detected a slight, though insignificant, increase in rubella virus antibody titer between timepoints, highlighting the potential loss of sensitivity due to removal of overlapping peptides. Insufficient serum prevented clinical testing for human herpesvirus 8 antibodies in this patient.

The conceptual paradigm underlying AVARDA may find applications beyond analysis of VirScan data. In any study involving library redundancy and potential antibody cross reactivity, any or all of the algorithm’s Modules may enable deconvolution of the data. For instance, libraries of bacterial, parasitic, or fungal pathogen peptidomes may be analyzed analogously to VirScan data. Screening IgE antibodies against libraries of allergen-derived peptides may present similar challenges due to the sharing of epitopes across related organisms^22^. Additionally, incorporating the conceptual framework of AVARDA into the design of peptide libraries may enhance their ultimate discriminatory power. Beyond peptide library screening, full-length protein library screening (e.g., via protein microarrays, PLATO, etc.) may also benefit from AVARDA’s consideration of shared sequences and potential cross-reactivity.

Furthermore, PhIP-Seq analysis of the human proteome can be performed in conjunction with VirScan. In the setting of undiagnosed encephalitis, for example, a combined human proteomic and pan-viral analysis could prove particularly informative. Bayesian interpretation of the joined data sets could potentially be performed under an assumption of *either* autoimmunity *or* viral infection (or bacterial infection, etc.), which may further improve diagnostic sensitivity. Incorporation of AVARDA into clinical diagnosis, and/or the interpretation of epidemiological studies, is therefore likely to become an important tool for linking human health with environmental exposures.

## Methods

### Serum Samples

#### DAISY Longitudinal Study

The <u>D</u>iabetes <u>A</u>uto Immunity <u>S</u>tudy in the <u>Y</u>oung (DAISY) is a prospective study of children with increased risk for Type 1 Diabetes. The details of the newborn screening^23^ and follow up^24^ have been previously published. Per protocol, serum was tested at 9, 15, and 24 months and, if autoantibody negative, annually thereafter; children found to be autoantibody positive were re-tested every 3-6 months. Recruitment took place between 1993 and 2004 and follow-up results are available through February 2018. Written, informed consent was obtained from the parents of study participants. The Colorado Multiple Institutional Review Board approved all study protocols.

#### SNIP Study

The <u>S</u>ingapore <u>N</u>eurologic Infections <u>P</u>rogram (SNIP) is a prospective cohort approved by the Singhealth Centralised Instituitional Review Board (CIRB Ref: 2013/374/E). The SNIP study aims to describe the epidemiology of CNS infections in Singapore; improve the diagnosis of etiologies of CNS infections through a systematic clinical, laboratory and neuroradiological evaluation and extensive diagnostic testing; evaluate the prognosis, long-term outcomes and socio-economic costs of CNS infections; and establish an archive of biological tissues from patients with encephalitis and CNS infections that can be utilized for future testing for emerging pathogens or non-infectious etiologies.

Subjects were enrolled from 6 hospitals in Singapore (Singapore General Hospital, Tan Tock Seng Hospital, National University Health System, Changi General Hospital, Khoo Teck Puat Hospital, Kandang Kerbau Hospital). Individuals, both male and female, more than one month of age, were enrolled upon admission to the hospital with clinical suspicion for CNS infections, or one or more of the following: Fever or history of fever (≥ 38°C) during the presenting illness; seizures; focal neurological deficits; CSF white cell count pleocytosis (> 4 WBC/uL); abnormal neuroimaging suggestive of CNS infection; abnormal electroencephalogram suggestive of CNS infection; depressed or altered level of consciousness, or no alternative etiology for acute paralysis identified. Patients with indwelling ventricular devices such as EVD and ventriculo-peritoneal shunts were excluded from the study. Blood samples were be obtained on enrollment into the study and then at 2 weeks after enrollment or at time of discharge.

#### VirScan Assay

VirScan screening was performed as described previously^5,6,25,26^. Briefly, we used a mid-copy T7 bacteriophage display library spanning the human virome, which consists of 96,099 56-aa peptide tiles that overlap adjacent tiles by 28-aa. BLT5403 *E. coli* (Novagen) was used to expand the library, which was then stored at -80°C in 10% DMSO. An ELISA was used to quantify IgG serum concentrations (using Southern Biotech capture and detection antibodies, cat# 2040-01 and 2042-05, respectively). Next, 2 μg of IgG was mixed with 1 mL of the VirScan library at a concentration of 1 × 10^10^ pfu (diluted in PBS) for each reaction. Following overnight end-over-end rotation of the phage and serum mixtures at 4°C, 40 μL of protein A/G coated magnetic beads (Invitrogen catalog numbers 10002D and 10004D) were added to each reaction (20 μL of A and 20 μL of G) which were rotated an additional 4 h at 4°C. Later the beads were washed three times and then resuspended in a Herculase II Polymerase (Agilent cat # 600679) PCR master mix using a Bravo (Agilent) liquid handling robot. This mix underwent 20 cycles of PCR. Subsequently, 2 μL of this amplified product underwent an additional 20 cycles of PCR, during which sample specific barcodes and P5/P7 Illumina sequencing adapters were added. This product was pooled and then sequenced using an Illumina HiSeq 2500 in rapid mode (50 cycles, single end reads).

#### Pairwise Differential Peptide Enrichment Analysis

Sequencing read counts for the VirScan peptides were normalized to the sample’s sequencing depth (nRCs), and a robust regression of the top 1,000, by abundance, Day 1 nRCs was used to calculate the ‘expected’ Day 14 nRCs. In addition to the samples, each 96 well plate contained 8 wells without antibody input (mock IPs) that served to provide expected background signals for single timepoint analyses. These samples were also used to calculate the expected standard deviations for peptides of a given nRC^13^. For pairwise analyses, the observed Day 14 nRCs minus the expected Day 14 nRCs for each peptide (the residuals) were normalized to their expected standard deviations, in order to calculate a ‘pairwise z-score’. The Day 1 versus Day 14 nRC scatter plots were generated in R. Two cases, Pt15 and Pt16, failed our quality control filter, due to poor Day 1 versus Day 14 correlations, and were therefore from further analysis.

#### Peptide-Virus Alignment Table

The peptide-virus alignment tables were created as follows. First, all viral genomes, including representative genomes that are in RefSeq and ‘neighbor’ strains that are not in RefSeq, were downloaded on May 2, 2017 in GenBank format from the NCBI Viral Genome Resource^27^. The host field of the GenBank files and the host column in the NCBI Viral Genome Resource neighbors file were then used to find viral strains that infect humans. Furthermore, all viruses annotated with human host in the Viral-Host Database (v170502) were included^28^. The human-host annotation was propagated from each viral strain to all strains of the same species. BLAST databases^29^ of nucleotide sequences of the human viruses were created using makeblastdb at sequence, organism and species levels. tblastn v2.2.31+ was run to create peptide-virus alignment tables (parameters: “-outfmt 6 -seg no -max_hsps 1 -soft_masking false -word_size 7 -max_target_seqs 100000”).

#### Network Analysis and Binomial Statistics

AVARDA was developed and implemented in Python 3.6, with dependencies on the packages NumPy, pandas, NetworkX, SciPy, and StatsModels. The software reads in a file of z-scores for each peptide and sample and outputs the list of significant viral infections, along with the associated p-values, assigned counts and peptides to each virus, and other relevant information used in the analysis. AVARDA can be downloaded for general use at github.com/LarmanLab/AVARDA, where further documentation is provided in a README.

##### Defining the maximal set of nonoverlapping antibody specificities

As described in the Module 1 section, a database of all seven amino acid overlaps between all peptides in the VirScan library is used to construct an undirected network graph, where each node represents an enriched peptide and each edge represents an overlap of at least a seven-amino acid stretch shared by the linked peptides. Our goal is to remove all connections within the graph by iterative deletion of peptides. The peptide(s) with the maximum connections (degree) in the graph are removed first and the degrees of remaining peptides recalculated. If there are multiple peptides of equivalent maximum degree within the graph that share an edge, they are ranked by their enrichment z-scores and the minimum z-scoring peptide is removed first. This process is repeated until the max degree of all peptides left in the graph is two. At this point, the only remaining graph structures are cycles (no end peptides), paths (two end peptides), and isolated single peptides. For cycles, the peptide with the lowest z-score is removed first, which results in a path. For paths, peptides are numbered starting at one end. Even numbered peptides are removed from paths with odd-numbered length. Paths of even-numbered length are split by removing the peptide with the lowest z-score, which results in a path of odd-numbered length and another path of even-numbered length. This process is iterated until no more edges remain.

##### Null probabilities for binomial testing

As described in Modules 2 and 3, when comparing sets of enriched peptides associated with each virus, we use binomial testing to determine the significance of the associations. The null model assumes the peptides aligning to each virus were randomly drawn from the VirScan library. For each virus with at least one enriched peptide alignment, we perform a binomial test using N = total number of enriched hits, k = hits aligning to virusi, and the null probability ‘f’, which is the total number of peptide alignments to virusi, divided by the total number of peptides in the VirScan library. All pairs of viruses that share enriched peptide alignments are evaluated as follows. First, a binomial test is performed on the non-shared (‘unique’) peptides, in order to weigh the evidence for each distinguishable infection on its own. For this test, f = (number of peptides aligning to virusi but not virusj) / (length of library – shared alignments between virusi and virusj). Similarly, N = number of enriched hits – shared virusi,j hits. A second binomial test is performed on the shared (indistinguishable) peptide alignments. Here, f = shared virusi,j alignments/(length of library – unique alignments to virusi – unique alignments to virusj). To determine significance, we set a p-value threshold of p < 0.01, and require that the total number of peptide alignments, k > 2. These parameters can be adjusted depending on the desired stringency of the analysis.

##### Viral comparisons, shared alignment removal, and indistinguishability tags

N is first initialized to the number of enriched, nonoverlapping peptides. All viruses are then ranked in order of significance based on binomial testing of each virus’s enriched peptide alignments. Next, the most significant virus is compared to every other virus which shares peptide alignments. In these comparisons, if either virus is found to have a statistically significant number of non-shared peptides while the other does not, the shared alignments are removed from the latter virus (case **i** in **Fig 3A**). If both viruses are found to have statistically significant numbers of non-shared peptides, the shared alignments remain associated with both viruses, thus contributing to the significance of both infections (case **ii** in **Fig 3A**). If neither virus is found to have a statistically significant number of non-shared peptides, binomial testing is performed on shared peptides to determine whether there is sufficient evidence for infection by at least one of the two viruses. If the viruses have a significant number of shared alignments, a numerical “indistinguishability tag” is annotated to the results to indicate that the algorithm was unable to accurately distinguish between infection by one or both of the viruses (case **iii** in **Fig 3A**). Once the pairwise comparisons with the most significant virus have been completed, this virus is then removed from consideration and N is decreased by the number of peptides that were uniquely assigned to that virus. This procedure of updating N thus removes peptides associated with significant infections from consideration in subsequent tests of peptides using the null model. The p-values for each remaining virus are recalculated, and the new top ranking virusi is selected for the next round of comparisons until all virusij pairs have been evaluated. Finally, all p-values of infection are adjusted for multiple hypothesis correction in order to consider multiple hypothesis testing for all infections. Ancillary relevant data are also printed to file.

#### Clinical Antibody Tests

All herpes simplex virus, varicella zoster virus, cytomegalovirus and Epstein-Barr virus antibody testing was carried out on a Diasorin Liason®, which uses chemiluminescent immunoassay technology, at the Johns Hopkins Pathology Core facilities. All other serological analyses were performed by Quest Diagnostics; human herpesvirus 6 and 8 antibodies were analyzed via immunofluorescence assay technology and human parainfluenza viruse 3 antibodies were analyzed via complement testing. Enterovirus testing was not done due the large volume of sera required per test. Due to the limited availability of serum, all samples were diluted 1:10 prior to testing. Final results were interpreted by a clinical pathologist.

## Author contributions

D.R.M., S.V.K., and H.B.L. conceived the project, designed the algorithm and wrote the manuscript. S.V.K. additionally implemented the algorithm in Python. T.Y. developed the software for analysis of Illumina sequencing data. F.B. developed the viral genome database and generated the virus-peptide alignment matrix. L-F.W., D.E.A., L.V., K.T. and W.C provided the encephalitis cohort samples, conducted sample selection, preliminary testing and provided useful insight in analyzing this cohort. K.K. provided valuable insight on statistical methods. M.C. interpreted all serological assay results. M.R. and K.W. provided the T1D longitudinal samples as well as valuable information on the patient’s vaccine history.

## Acknowledgements

This work was made possible by support from the NIAID U24 Award AI118633 and the US Army Research Office Award W911NF-1410490. The DAISY team is supported by funding from the NIDDK R01 award DK032493. We acknowledge the Singapore Infectious Diseases Initiative (SIDI) for funding support and would like to thank the study team and colleagues of all the hospitals involved. The Duke-NUS team is supported by funding from NMRC (NMRC/ZRRF/0002/2016) and CDPHRG/0006/2014) and NRF (NRF2016NRF- NSFC002-013).

---

**Supplemental Figure 1.**
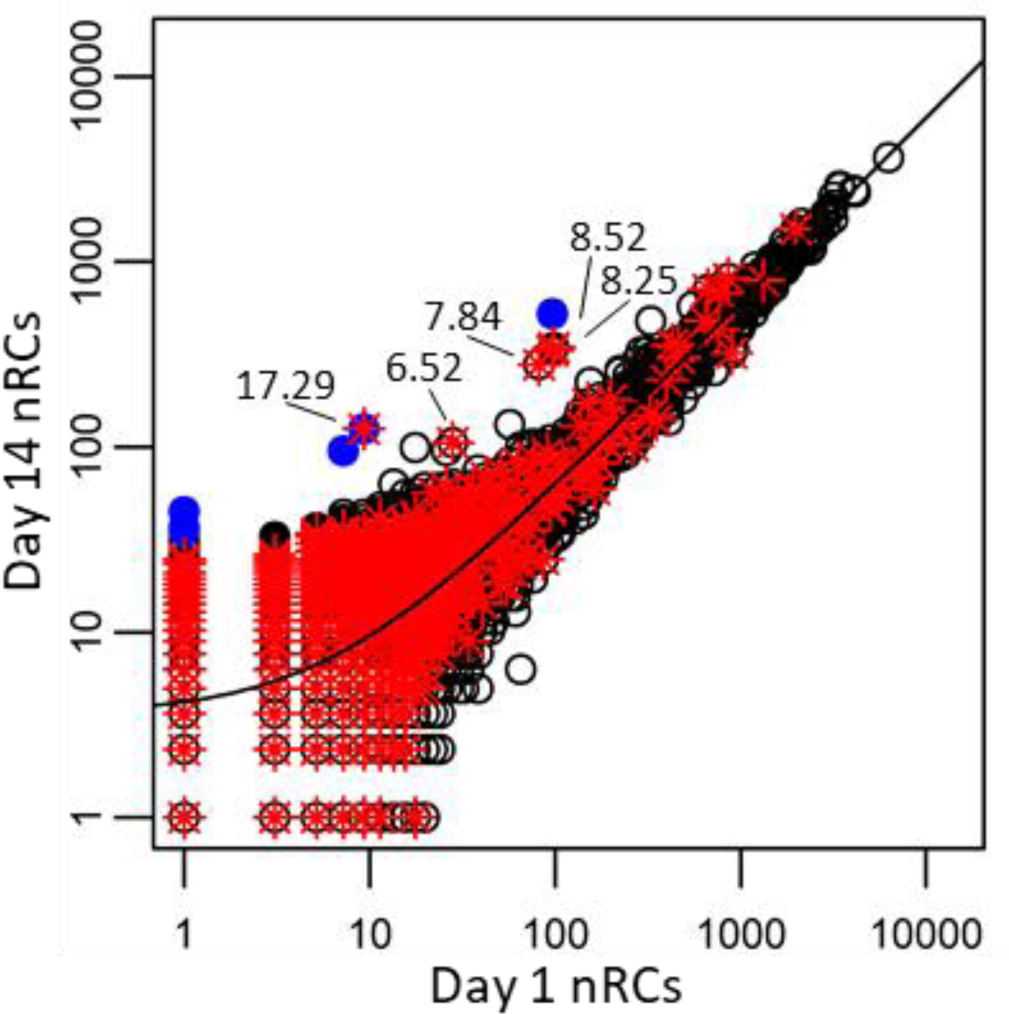
Dependence of sensitivity on enrichment threshold. The scatterplot shows Pt9’s VirScan normalized read counts from the day 1 dataset (x-axis) plotted against the day 14 dataset (y-axis). A robust linear regression line is shown. Points in blue represent peptides enriched with pairwise z-scores >10. Peptides with alignments to varicella-zoster virus (VZV) are additionally marked with a red star. The top 5 differentially enriched VZV peptides are additionally annotated with their pairwise enrichment scores.

**Supplemental Figure 2.**
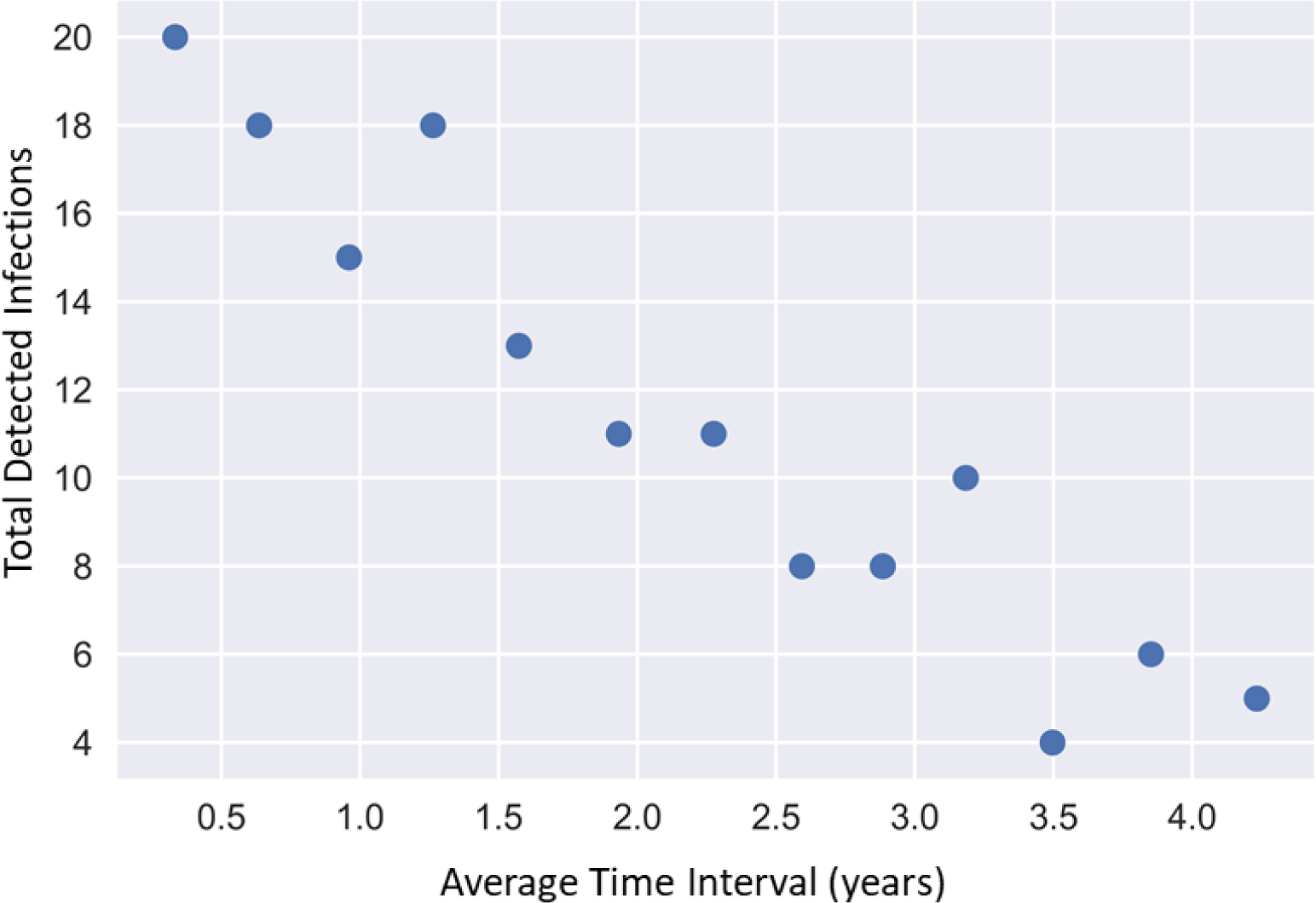
Scatterplot of sampling frequency versus number of significant infections identified by AVARDA. The horizontal axis corresponds to the average time (in years) between patient serum samples used for AVARDA analyses. The vertical axis corresponds to the cumulative number of viral infections identified over the time course.

**Supplemental Figure 3.**
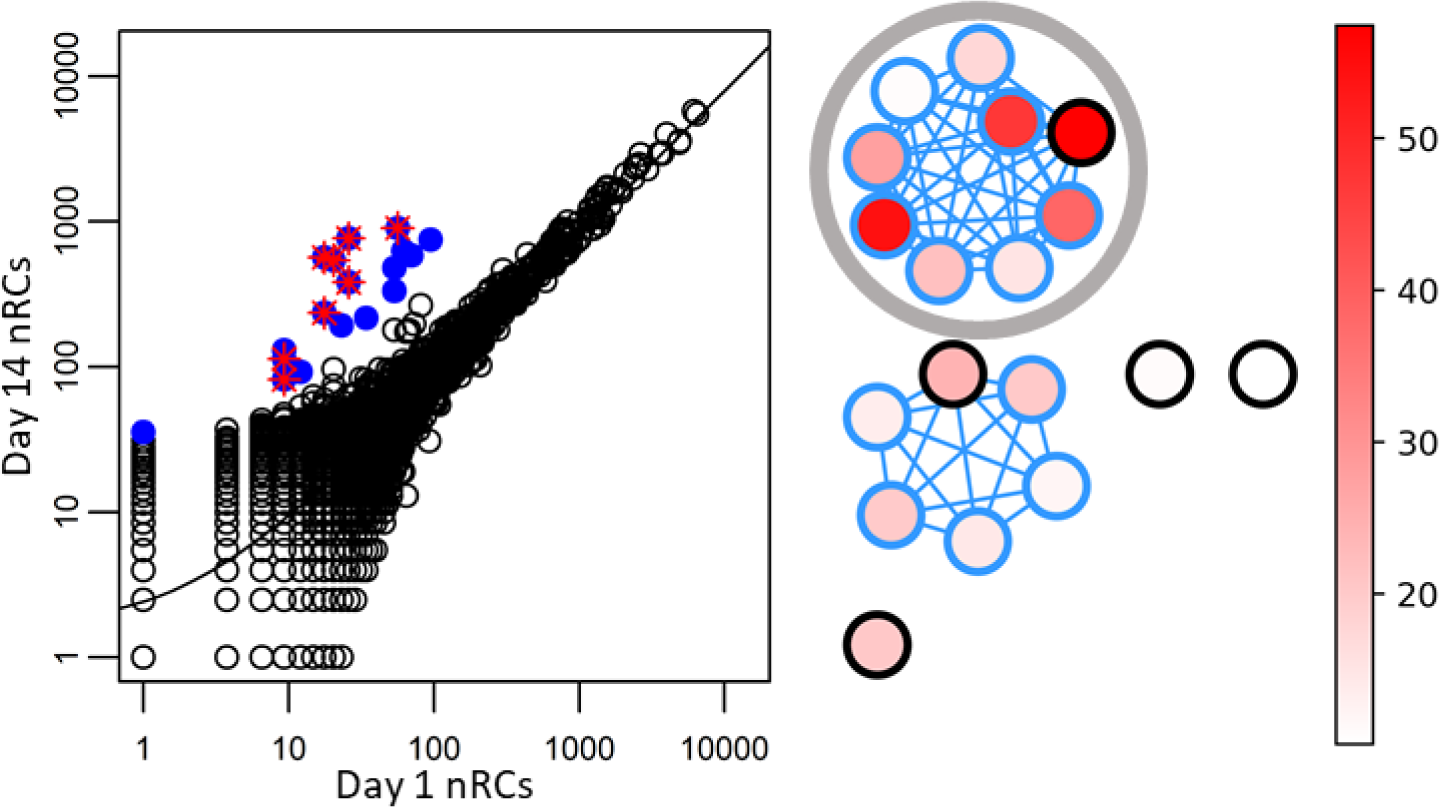
Impact of network filter on AVARDA. **(A)** Pairwise scatterplot of Pt17’s normalized read counts (nRCs). Points in blue represent enriched peptides. Enriched peptides associated with rubella are additionally marked in red. **(B)** Network graph representation of seven amino acid overlaps among the pairwise enriched peptides of A. Peptides in the circled sub-cluster align with Rubella virus, peptides in the other cluster align with human herpesvirus 8. Peptides are colored according to z-score and outlined in black if selected by the independence filter.

**Supplemental Table 1.** Pairwise enriched VirScan peptides for Pt1. Each row indicates the number of pairwise-enriched peptides designed to represent each virus.

| Represented Virus | Number of Enriched Peptides |
| --- | --- |
| Human herpesvirus 1 | 68 |
| Human herpesvirus 2 | 22 |
| Cercopithecine herpesvirus 16 | 14 |
| Varicella-zoster virus | 11 |
| Cercopithecine herpesvirus 1 | 10 |
| Herpes simplex virus type 2 | 3 |
| Human cytomegalovirus | 3 |
| Influenza A virus | 3 |
| Human enterovirus 71 | 2 |
| Sandfly fever sicilian virus | 2 |
| Xenotropic MuLV-related virus | 2 |
| Yellow fever virus | 2 |
| Banna virus | 1 |
| Epstein-Barr virus | 1 |
| Guanarito mammarenavirus | 1 |
| Hepatitis C virus genotype 1c | 1 |
| Hepatitis delta virus | 1 |
| Human adenovirus 14 | 1 |
| Human adenovirus C serotype 1 | 1 |
| Human immunodeficiency virus type 1 | 1 |
| Human papillomavirus type 29 | 1 |
| Human papillomavirus type 37 | 1 |
| Human respiratory syncytial virus B | 1 |
| Orf virus | 1 |
| Pichinde mammarenavirus | 1 |
| Pseudocowpox virus | 1 |
| Punta toro phlebovirus | 1 |
| Sabia mammarenavirus | 1 |
| Vesicular stomatitis Indiana virus | 1 |

